# Spatial and Spectral Changes in Cortical Surface Potentials during Pinching versus Thumb and Index Finger Flexion

**DOI:** 10.1101/2024.09.30.615538

**Authors:** Panagiotis Kerezoudis, Michael A Jensen, Harvey Huang, Jeffrey G. Ojemann, Bryan T. Klassen, Nuri F. Ince, Dora Hermes, Kai J Miller

## Abstract

Electrocorticographic (ECoG) signals provide high-fidelity representations of sensorimotor cortex activation during contralateral hand movements. Understanding the relationship between independent and coordinated finger movements along with their corresponding ECoG signals is crucial for precise brain mapping and neural prosthetic development. We analyzed subdural ECoG signals from three adult epilepsy patients with subdural electrode arrays implanted for seizure foci identification. Patients performed a cue-based task consisting of thumb flexion, index finger flexion or a pinching movement of both fingers together. Broadband power changes were estimated using principal component analysis of the power spectrum. All patients showed significant increases in broadband power during each movement compared to rest. We created topological maps for each movement type on brain renderings and quantified spatial overlap between movement types using a resampling metric. Pinching exhibited the highest spatial overlap with index flexion, followed by superimposed index and thumb flexion, with the least overlap observed for thumb flexion alone. This analysis provides practical insights into the complex overlap of finger representations in the motor cortex during various movement types, and may help guide more nuanced approaches to brain-computer interfaces and neural prosthetics.

**SIGNIFICANCE STATEMENT:** This study measured brain activity actively during finger movement and found that pinching movements show higher cortical overlap with index finger flexion and uniquely engage parietal regions, advancing our understanding of motor control hierarchy and sensorimotor integration, while suggesting improvements for more naturalistic brain-computer interfaces through prioritized index finger decoding and integration of parietal lobe measurements.

## INTRODUCTION

The hallmark of human motor behavior is the ability to execute finely coordinated individual finger movements. Dextrous hand gestures are crucial for performing daily tasks and can be considered as a combination of individual finger movements with varying degrees of freedom, particularly in flexion and extension.[1] Interestingly, certain movements, including reaching, grasping (palmar or pincer), carrying (transporting), and placing have been previously proposed as motor primitives.[2,3] These primitives represent the building blocks for executing goal-directed actions, such as a monkey reaching for a seed, grasping it, bringing it towards the mouth and finally eating it.[2]

Cortical representations of individual and composite finger movements are an area of ongoing investigation in electrophysiology and neuroscience.[1,4] Specifically, brain-computer interface (BCI) research has made significant progress in examining cortical dynamics related to motor function and decoding individual finger movements. Studies using electrocorticography (ECoG) and intracortical recordings have demonstrated the ability to distinguish between movements of different fingers with accuracy that can exceed 80%.[5–8]

However, the neural representation of compound movements and motor primitives with individual fingers has not yet been fully elucidated. While some studies suggest that complex hand movements might be represented as a combination of individual finger movement patterns[9,10], others indicate that there might be distinct neural signatures for coordinated actions like reaching and grasping.[11,12] It remains unclear to what extent convergent hand movements represent sums of the individual finger movements or activate different brain areas, such as the premotor cortex and posterior parietal cortex. This area of research is particularly relevant for the development of more naturalistic neuroprosthetic control systems.

To address this knowledge gap, we examined ECoG recordings from three patients with medically refractory epilepsy undergoing seizure workup. We analyzed the spectral and spatial changes during thumb flexion, index flexion, and pinching, quantifying the spatial overlap between these movement types. Our study aims to provide insights into the neural representations of individual and composite finger movements, with implications for BCI technology and prosthetic control. Understanding the relationship between individual finger movements and composite actions like pinching could inform more sophisticated decoding algorithms, potentially enabling more intuitive and dexterous control of neuroprosthetic devices.

## MATERIALS AND METHODS

### Ethics statement

All patients participated in a purely voluntary manner, after providing informed written consent, under experimental protocols approved by the Institutional Review Board of the University of Washington (#12193). All patient data was anonymized according to IRB protocol, in accordance with HIPAA mandate. It was made available through the library described in “A Library of Human Electrocorticographic Data and Analyses”, freely available at https://searchworks.stanford.edu/view/zk881ps0522.[13] No analyses of these data have been previously published.

### Patient population

We recruited 3 patients (2 females, aged 18, 21 and 19 years) that underwent invasive clinical EEG monitoring for localization of seizure foci as part of their workup for medically refractory epilepsy using subdural ECoG grids. All patients were right-handed.

### Electrical recordings

The platinum ECoG arrays were configured as a combination of grid (8×8 or 4×8) arrays and strip arrays, numbering a total of 32 to 81 contacts (i.e. channels). The diameter of the electrode contacts was 4mm and the inter-electrode distance was 10mm. These arrays were surgically placed by one of the senior authors (J.G.O). The ECoG signals were split into two identical sets, one towards the clinical EEG system (XLTEK, Oakville, Ontario, Canada) and the other to a research system (Synamps2, Neuroscan, El Paso, TX), which were biosignal amplifiers at 1 kHz with a bandpass-filter at 0.3-200 Hz. Finger position was recorded using a piezoelectric sensor dataglove (5DT, Irvine, CA).[14] The general purpose BCI2000 software was used for stimulus presentation and ECoG signal data collection.[15]

### Cortical rendering and electrode localization

The cortical surface from a preoperative MRI was rendered using either Freesurfer or Spm5 software in order to determine the relationship between the gyral anatomy and electrode position.[14,16] The electrode positions were calculated after co-registering the post-operative computed tomography (CT) to the pre-operative MRI using the CTMR package by Hermes et al, 2010, which has been shown to accurately localize the electrode positions within a ∼4mm error.[17]

### Movement tasks

Subjects were presented with a word cue displayed on a bedside monitor to perform the following self-paced movement tasks, each on a separate run: 1) pinching move between the thumb and the index finger or 2) individual thumb or index finger flexion. A 2-second rest trial (blank screen) followed each movement trial. There were up to 30 cues for each movement type. Segmentation of movement vs rest period was performed based on visual inspection of the data glove trace and manually marking the initiation and termination of movement.

### Signal processing

#### Pre-processing

The potential measured at each electrode was re-referenced with respect to the common average of all electrodes. Electrodes were visually inspected for significant artifact or epileptiform activity, removed from re-referencing & omitted prior to further analysis.

#### Power spectral snapshots

Power spectral snapshots using Welch’s method with 1s Hann window were generated for the entire experiment in each electrode as well as separately for movement and rest blocks. In addition, we mean-normalized the log power spectra by dividing the power spectrum for each trial with the mean electrode power spectrum.

#### Dynamic power spectrum

We generated time-frequency approximations after convolving the voltage time-series data with a Morlet wavelet (10 cycles) in order to estimate the amplitude and phase of the signal across the frequency range for every point in time.

#### Power spectrum decoupling

We performed decoupling of the power spectra using a previously established method by Miller et al, 2009.[18] In summary, we performed Principal Component Analysis (i.e. eigendecomposition) of the power spectral density-covariance matrices reveal which frequencies vary in power together and are ordered according to the value of the corresponding eigenvalue. It has been shown that the 1^st^ PSC corresponds to broadband spectral changes (i.e. spanning all frequencies), whereas the 2^nd^-4^th^ PSCs typically capture rhythmic power spectral phenomena (i.e. mu rhythm, alpha rhythm etc).[18] In addition, the projection of the dynamic spectrum onto the 1^st^ PSC will yield the logarithm of the time course of the power spectrum power law coefficient P(f, t) = A(t)\**f*^−χ^.[19] This logarithm was subsequently smoothed with a Gaussian window (SD 250 ms), z-scored and re-exponentiated in order to obtain the broadband traces plotted in **Figure 1**.

**Figure 1.**
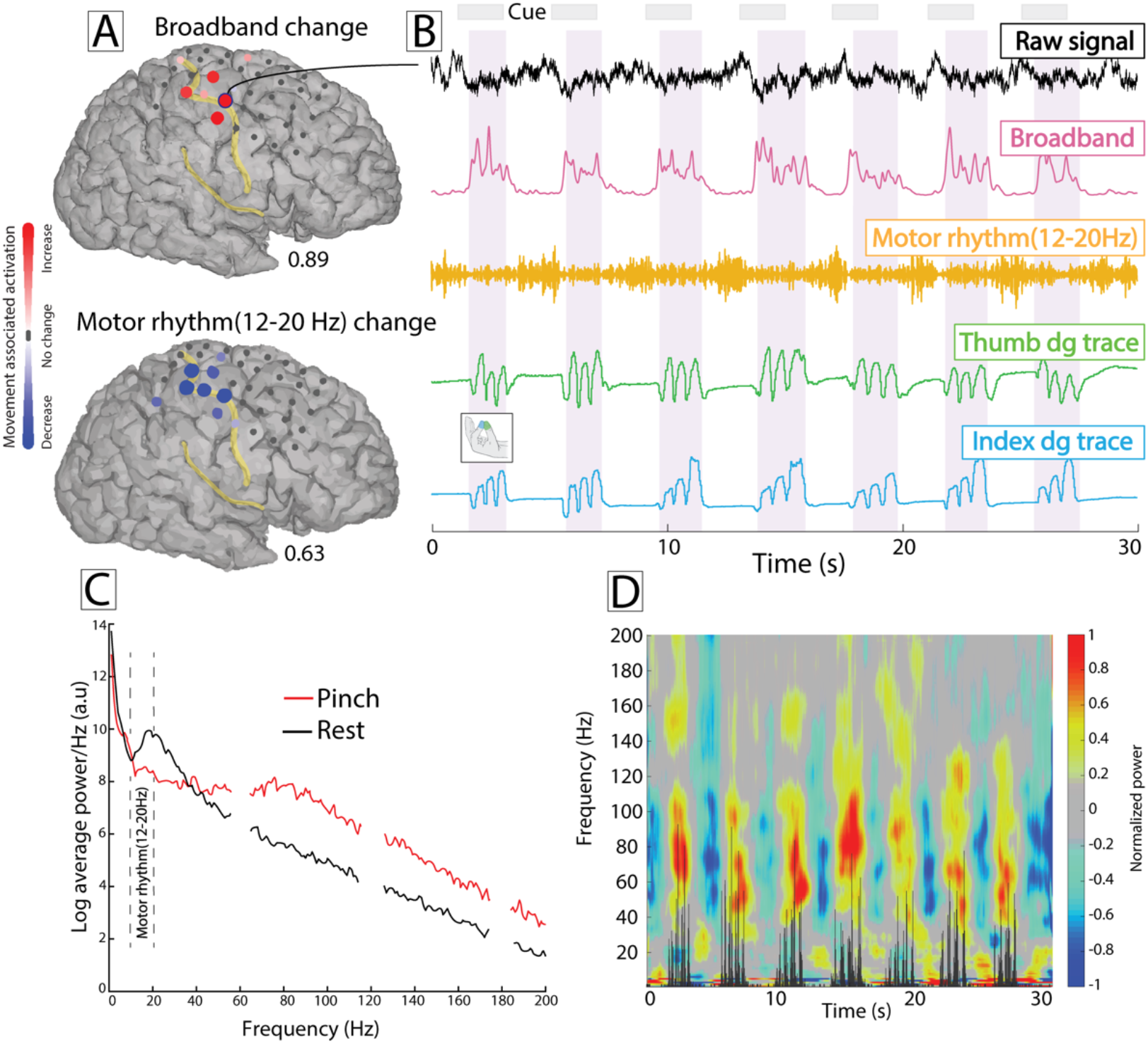
Electrocorticographic (ECoG) activity in the right frontoparietal cortex during pinch movement and rest (subject 1). **(A)** Topological maps demonstrating focal broadband power increase (color represents signed r^2^ measurement, scaled to the maximum across the array, i.e. 0.89) and more widespread spatial distribution in motor rhythm (12-20 Hz) power decrease (with Bonferroni correction). **(B)** The raw ECoG voltage, broadband and motor rhythm power time series from in the precentral gyrus during 30 s of recording are shown. Thumb and index movement as captured by the data glove are shown at the bottom panels. The gray boxes represent the screen cues, while the light purple shaded regions represent the movement epochs. Broadband spectral changes demonstrate excellent correlation with finger movement. **(C)** Power spectral density plot from the entire behavioral task (same channel) showing decrease in low-frequency oscillations and broadband increase across the rest of the frequency range, during movement (red) compared to rest (black). **(D)** Time-frequency spectrogram during the same period of recording. The black trace inset represents the rectified 1st derivative of thumb movement.

This broadband power time course has been shown to be a robust estimate of behaviorally relevant local cortical activity as well as predictive of the finger movement time course, as measured by the data glove.[20] Finally, we calculated the signed r^2^ cross-correlation values from broadband activity, between each movement modality (pinching, thumb flexion, or finger flexion) and rest. Movement trials were compared with the rest trials that followed the same movement type.[21] Thus, r^2^ represents the percentage in variance in the joint distribution of movement-rest blocks that can be explained by a difference in the movement and rest trial means, respectively:

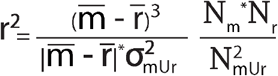

#### Quantifying spatial overlap

The spatial extent of overlap and degree of change in broadband power activity was quantified and compared using a resampling metric, as described previously.[13] In summary, for measures of type X_n_ and Y_n_ (n denoting the electrode), the true overlap (O^T^_XY_) was calculated from the pairwise dot product of the broadband r^2^ value. The spatial overlap metric was calculated by dividing the true overlap with the maximum possible overlap (O^M^_XY_) by computing the dot product of distributions X_n_ and Y_n_ assorted in descending order. The electrodes were subsequently scrambled 10^6^ times (‘n’→’m’) and a surrogate overlap was obtained from each permutation (O^S^_XY_) Finally, the p-value for statistical significance was estimated as the percentage of surrogate distributions O^S^_XY_ that were greater than O^T^_XY_ (or the percentage that were less in case O^T^_XY_ < 0) divided by the total number of shuffles (10^6^). The methodology is graphically displayed in **Supplemental Figure 1**. The r^2^ electrode activation maps of the following conditions were compared with pinching [summarized in **Table 1**]: finger flexion, thumb flexion, their maximum value per channel, their geometric mean and modified geometric mean.

**Table 1.** Description of the individual and composite metrics used to compare spatial overlap between pinching and finger flexion. Each r^2^ value corresponds to a single channel.

| Condition | Formula | Rationale |
| --- | --- | --- |
| Thumb flexion | $r_{thb}^2$ | Components of pinching movement. |
| Index flexion | $r_{idx}^2$ | |
| Maximum value of thumb flex,<br>index flex | $\max\{r_{thb}^2, r_{idx}^2\}$ | Channel activation reflects the maximal activity evoked by each individual finger movement (i.e. the union of two activation sets). |
| Geometric mean of thumb flex and<br>index flex | $\sqrt{\vert r_{thb}^2 * r_{idx}^2\vert }$ | Channel activation reflects the average activity evoked by each individual finger movement (i.e the intersection of two activation sets). |
| Modified geometric mean of<br>thumb flex and index flex | $1 - \sqrt{\vert (1 - r_{thb}^2) * (1 - r_{idx}^2)\vert }$ | Channel activation reflects the complement of the average activity not explained by each individual finger movement (i.e the complement of the intersection of the non-activation sets). |

**Abbreviations:** idx: index; thb: thumb

#### Broadband activation latency

For subject 3, we determined the time of onset of significant broadband activity for each channel using a previously described robust estimate on a single-trial level, by Foster et al.[22] In summary, the procedure is: First, we extracted the screen cue and movement onset time indices for each movement trial, and we selected a window 0.5 s prior - 3 s following the screen cue onset. Next, contiguous time points of broadband activity, at least 100 ms in duration, above 2 standard deviations of the baseline (i.e. first 500 ms of the window) were identified. At the first time point above the threshold, a 200 ms window was extracted which was further divided into 10 segments of 100 ms with 90% overlap. Linear regression was subsequently performed on each of the 20 segments to obtain slope and residual error. The top five segments with the largest slope coefficients were selected, and the segment with the least mean squared error was defined as the “onset” segment. The difference between the first time point of the onset segment and movement onset defined the latency to broadband activation for the single trial. For each channel, we averaged the latencies across all trials using the median.

### Code availability

All code necessary to reproduce findings will be made publicly available on GitHub following the publication of the manuscript. All analysis was conducted in MATLAB (R2022b, Mathworks, Natick, MA, USA). Figures were produced in MATLAB and finally edited in Adobe Illustrator 2024 (Adobe, San Jose, CA, USA).

## RESULTS

### Spectral changes related to movement

Across all 3 subjects, we observed significant increase in broadband power with movement and decrease in power in narrowband, low-frequency oscillations [**Figure 1A-D**]. Specifically, the maximum channel r^2^ values associated with movement were 0.63/0.72/0.89 for thumb flexion, 0.83/0.67/0.66 for index finger flexion and 0.89/0.80/0.64 for pinching. Broadband power increases following each movement type were more spatially focused, whereas motor beta rhythm (12-20 Hz) power decrease was more widely distributed over the cortical surface. We found that the time course of broadband activation matched finger movement with high fidelity [**Figure 1B**].

### Somatotopy of movement-related spectral changes

The r^2^ activation values for each channel in the subdural grid were plotted on the 3D brain renderings for each individual finger movement type as well as their composite metrics **[Figure 2]**. Comodulation between thumb and index flexion was associated with significant overlap by 50%-68% in r^2^ activation. Furthermore, in all three patients, spatial overlap between pinch and index flexion (69-87%, all p<.001) was considerably higher compared to pinch and thumb flexion (27-77%; significant in 2 of 3 patients). The modified geometric mean metric performed similarly or better compared to the index flex (61-96%), while the max (39-89%) and geometric mean (68-81%) metrics ranked in between.

**Figure 2.**
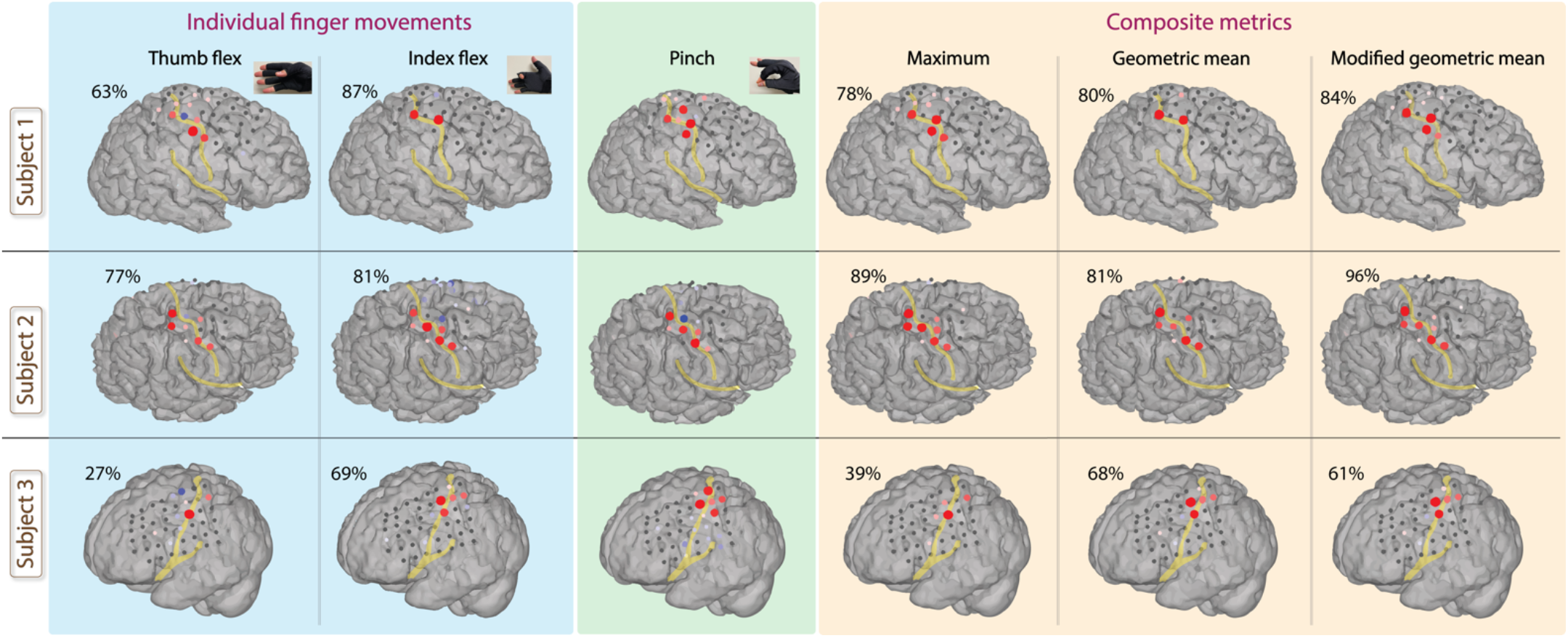
Topological maps in each subject demonstrating changes in broadband power at different cortical sites for thumb flexion, index finger flexion (blue panel), pinching (green panel) and three r^2^ metrics combining thumb and index movement (orange panel): by taking the maximum r^2^ across the two finger movements, the geometric mean and the modified geometric mean (see Methods for detailed description). The spatial overlap of each metric with pinching is shown on the top left corner of the brain renderings. Overall, it can be appreciated that index flexion, followed by the modified geometric mean, had the highest overlap with pinching.

In subject 3, we observed activated channels in the postcentral gyrus during pinching that were (relatively) silent during either individual finger movement. Based on response onset latency analysis, these channels either preceded M1 activation by approx. 80 ms, or anteceded by 20-100 ms **[Figure 3]**.

**Figure 3.**
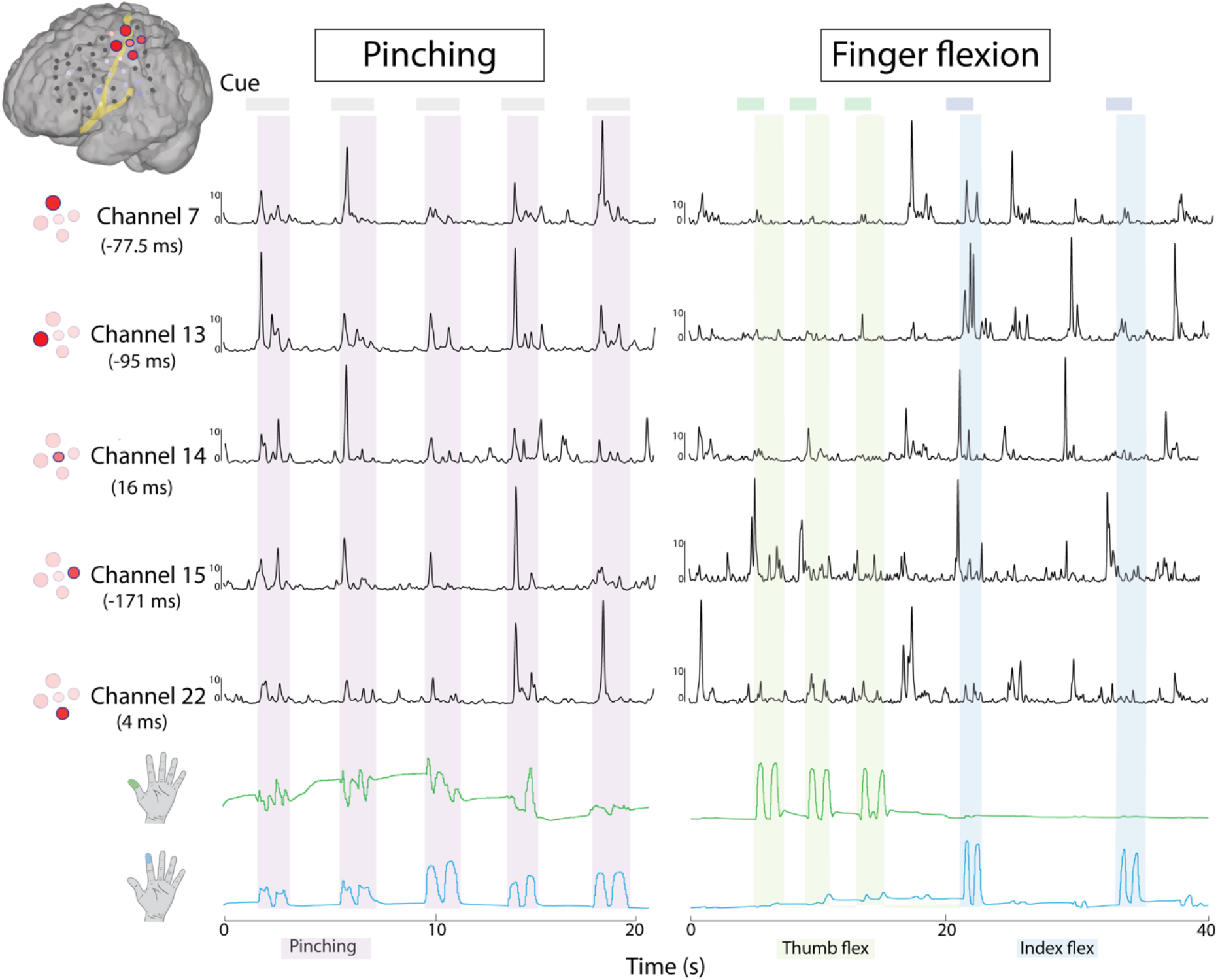
Broadband timeseries across most significant channels during pinching (light purple), thumb (light green) and index finger (light blue) flexion trials (subject 3). The number in parentheses below the channel number corresponds to the median broadband latency (please refer to Methods for more details).

## DISCUSSION

In this study, we analyzed cortical activity in three patients with subdural ECoG arrays during thumb flexion, index flexion, and pinching movements. Our findings reveal important insights into the neural representations of these movements and their implications for brain-computer interface (BCI) design. We demonstrated that the cortical representation of pinching partially overlaps with individual finger movements, particularly the index finger, and can involve postcentral areas as well. Interestingly, our nonlinear composite metrics did not lead to significantly higher overlap compared to index finger representation alone, suggesting a complex relationship between individual and composite movements in cortical encoding.

Our study revealed a higher spatial overlap between pinching and index finger flexion neural representations compared to thumb flexion, particularly in the precentral gyrus. Note that this observation cannot be explained by performing a pinch in which the index finger moves while the thumb is held still since it holds up when the thumb is clearly moving during pinch movement (Figures 1 and 3). This overlap has important implications for prosthetic device control, especially in scenarios with limited channel numbers or suboptimal electrode placement. For prosthetic devices with limited degrees of freedom or computational power, focusing on decoding index finger movements might provide the most efficient use of resources.[23,24] Our results suggest that accurate index finger control could potentially enable a range of composite movements, including pinching, without the need for separate thumb control. In a similar fashion, in clinical settings where perfect electrode placement is not always possible, training users to control index finger movements might naturally translate to improved control of composite movements like pinching.[25]

The use of clinical ECoG grids in our study, while not as high-density as micro-electrode arrays, provided us with the unique advantage of capturing neural activity across a wider cortical area. This broader coverage allowed us to simultaneously record from both the precentral (motor) and postcentral (somatosensory) gyri. The ability to capture this expansive neural landscape proved crucial in identifying the differential activation patterns between pinching and individual finger movements, particularly the parietal involvement in pinching.[26] Notably, in one patient, we observed activation in parietal areas during pinching that was not present during individual index or thumb flexion. This finding underscores the critical role of sensory feedback in the neural representation of pinching movements. The parietal cortex, known for its involvement in sensorimotor integration, appears to play an independent specific role in the complex task of pinching, possibly due to the increased sensory demands of finger-to-finger contact.[27,28]

Therefore, while focusing on index finger decoding from the motor cortex may provide a good foundation for prosthetic control, our findings regarding parietal activation during pinching suggest that relying solely on precentral gyrus motor cortex signals may not be sufficient for fully natural pinching movements. Future BCI designs might consider incorporating signals from both motor and somatosensory areas to capture the full complexity of intended gestures.[29] Integrating signals from parietal regions could significantly enhance the “naturalness” and precision of pinching movements in neuroprosthetic applications. This approach could lead to adaptive BCI systems that can switch between low-level individual finger control and high-level composite movement modes based on the detected spatial patterns of activation across both motor and sensory cortices.[30]

### Limitations

This work is limited by the small cohort size and the use of standard-scale clinical ECoG, rather than high-density research grids, the latter have been shown to have superior spatial performance for 6 elementary movements in the alpha, beta and gamma frequency bands.[8] We were unable to incorporate kinetics and/or kinematics in our analysis to correlate with the ECoG signal. The dataglove captures mean deformation of an electrically conductive elastomer with piezoresistive properties, but does not allow for detailed analysis of force of contraction or degree of displacement/range of motion in the interphalangeal joints. Finally, we elected not to perform decoding, as the primary aim of this work was inference of cortical topology and not prediction of finger kinematics.

## CONCLUSION

Our analysis revealed that the ECoG signal for pinching movements comprises a combination of signals from individual thumb and index movements, with a considerably larger overlap with the latter. Our findings not only provide valuable insight into the overlap of finger representations in the motor cortex during different actions, but also underscore the importance of integrating sensory mechanisms (whether actually experienced or only anticipated) and parietal signals for achieving more natural and precise control, especially for complex movements like gestures. Future neuroprosthetic designs might leverage the observed overlap patterns and the unique neural signatures of composite movements as independent features. This approach has the potential to refine currently available BCI control algorithms, enhance their therapeutic efficacy, and ultimately provide users with more comprehensive and intuitive control of prosthetic devices.

---

**Supplemental Figure 1.**
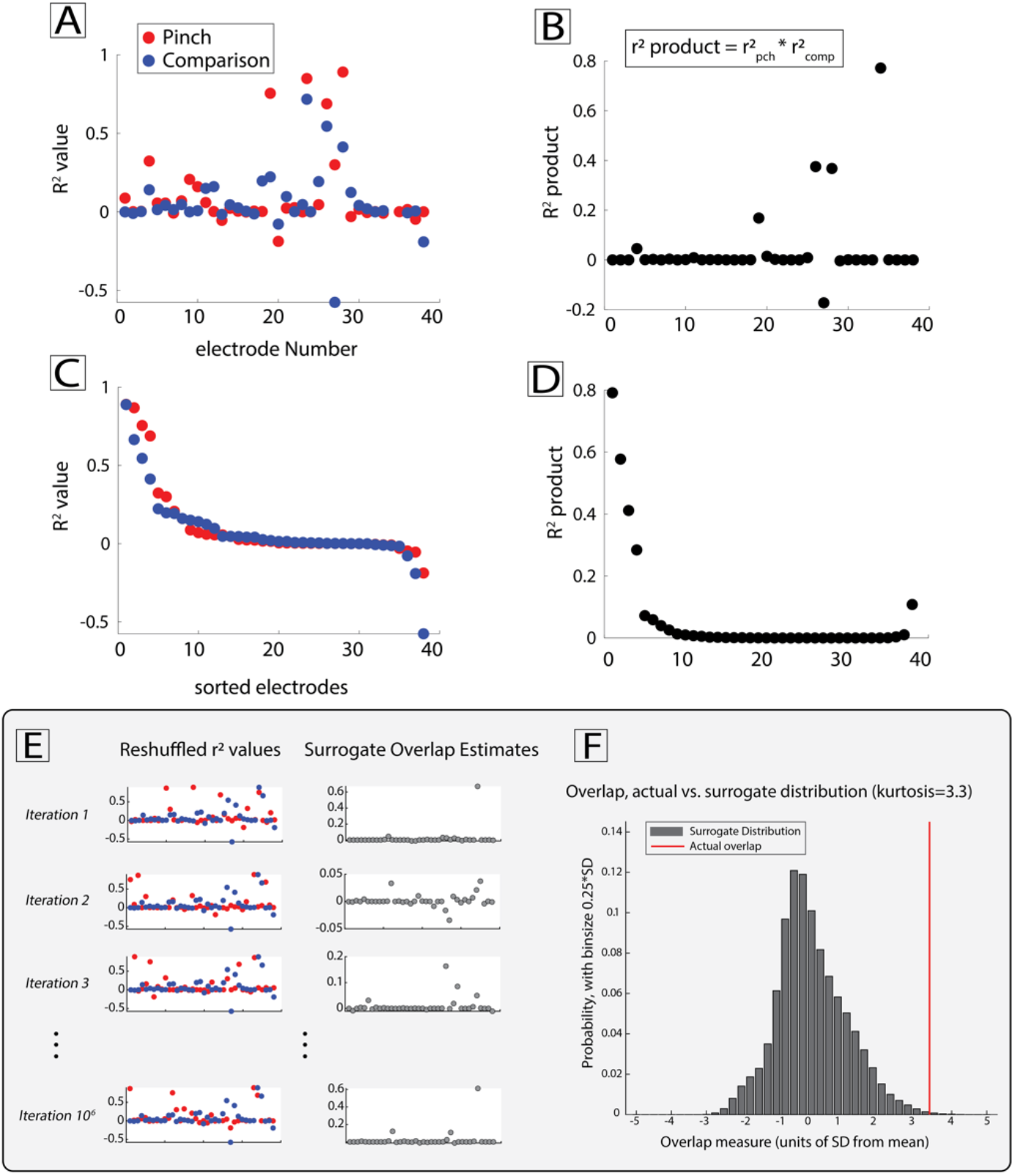
Demonstration of the method for quantifying the overlap in spatial distribution. Example data are shown from subject 1. **(A)** The r^2^ value, as a metric of activation during movement vs rest, is computed and plotted for each electrode, during pinching versus thumb flexion, index flexion or one of the other metrics we used (collectively labeled here as ‘comparison’). **(B)** The overlap can be quantified as the dot product across electrodes, thereby denoting that activities are paired together at each electrode. **(C)** Electrodes are sorted in decreasing r^2^ order and **(D)** the dot product is recalculated, corresponding to the maximum possible overlap. **(E)** We can probe all of the possible configurations of activation values by reshuffling the electrode positions (left panels) for one of the conditions (pinch) and recalculating the dot products (right panels). This is done 106 times, generating the distribution of surrogate dot products. **(F)** The generated surrogate distribution can be used to estimate a p-value for the significance of the overlap; in this case the distribution tends to be approximately Gaussian (kurtosis = 3.3). P-value is defined as the area to the right of the actual value (red line) to the total area under the curve of the distribution.

